# Short-term elevations in glucocorticoids do not alter telomere lengths: A systematic review and meta-analysis of non-primate vertebrate studies

**DOI:** 10.1101/2021.05.19.444844

**Authors:** Lauren Zane, David C. Ensminger, José Pablo Vázquez-Medina

## Abstract

**Background:** The neuroendocrine stress response allows vertebrates to cope with stressors via the activation of the Hypothalamic-Pituitary-Adrenal (HPA) axis, which ultimately results in the secretion of glucocorticoids (CORT). Glucocorticoids have pleiotropic effects on behavior and physiology, and might influence telomere length dynamics. During a stress event, CORT mobilizes energy towards survival mechanisms rather than to telomere maintenance. Additionally, reactive oxidative species produced in response to increased CORT levels can damage telomeres, also leading to telomere shortening. In our systematic review and meta-analysis, we tested whether CORT levels impact telomere length and if this relationship differs among time frame, life history stage, or stressor type. We hypothesized that elevated CORT levels are linked to a decrease in telomere length.

**Methods:** To test this hypothesis, we conducted a literature search for studies investigating the relationship between telomere length and CORT levels in non-human vertebrates using four search engines: Web of Science, Google Scholar, Pubmed and Scopus, last searched on September 27th, 2020. This review identified 31 studies examining the relationship between CORT and telomere length. We pooled the data using Fisher’s Z for 15 of these studies. All quantitative studies underwent a risk of bias assessment. This systematic review study was registered in the Open Science Framework Registry (https://osf.io/rqve6).

**Results:** The pooled effect size from fifteen studies and 1066 study organisms shows no relationship between CORT and telomere length ((Fisher’s Z= 0.1042, 95% CI = 0.0235; 0.1836). While these results support some previous findings, other studies have found a direct relationship between CORT and telomere dynamics, suggesting underlying mechanisms or concepts that are not currently taken into account in our analysis. The risk of bias assessment revealed an overall low risk of bias with occasional instances of bias from missing outcome data or bias in the reported result.

**Conclusion:** We highlight the need for more targeted experiments to understand how conditions, such as experimental timeframes, stressor(s), and stressor magnitudes can drive a relationship between the neuroendocrine stress response and telomere length.

## Introduction

The vertebrate neuroendocrine stress response integrates external stimuli into a broad range of physiological adjustments through the activation of the Hypothalamic-Pituitary-Adrenal axis (HPA-axis) and the concomitant secretion of glucocorticoids (CORT) [1–2]. While the primary CORT produced varies by taxa (e.g., cortisol in humans and corticosterone in birds and other mammals [3]), the impacts of CORT on organismal physiology are remarkably similar. Across species, an increase in CORT secretion can typically be detected in 3-5 minutes following interaction with a stressor [4]. Additionally, CORT are relatively easy to quantify because they are present in all vertebrates and can be measured noninvasively in multiple matrices including hair and feces using a variety of assays [5–6]. Therefore, wildlife stress physiology studies often rely on CORT measurements as a stress indicator [7]. Following their secretion, CORT induce a myriad of acute behavioral and physiological effects to prioritize immediate survival [8–9].

In addition to allowing animals to cope with immediate stressors, CORT influence other cellular processes such as telomere length dynamics. Telomeres are evolutionarily conserved caps that protect chromosomes against the loss of coding nucleotides during cell replication and against chromosomal fusions [10]. Telomere shortening is associated with aging, stress, and survival, and is thus of interest to several fields of biology [1,11]. In humans, increased telomere loss predicts the onset of age-related diseases such as cardiovascular complications, cellular senescence, and other aging phenotypes [12–13]. Telomere attrition can be attributed to several causes including the “end replication problem” in which the terminal end of linear DNA cannot be completely replicated by the lagging strand [14]. Since the end replication problem occurs at every cell division, telomeres continuously shorten with age progression [15]. Other stressors such as inflammatory challenges erode telomeres regardless of age [16].

In non-human vertebrates including birds, mammals, fish, amphibians and reptiles, exposure to challenging environmental conditions correlate with shorter telomeres [17–18]. Reproductive stressors such as an artificially increased brood size can also shorten telomeres in zebra finch parents compared to controls and parents with a reduced brood size [19]. Early telomere length is positively correlated to survival and lifetime breeding success in both wild purple-crowned fairy wrens and zebra finch. Thus individuals with longer telomeres are more likely to survive and produce more offspring that survive to maturity [20–21]. Therefore, telomere dynamics--the change in telomere length attributed to processes of elongation and shortening--is related to organismal fitness [22]. In addition to impacting telomere length, stressors that lead to energy limitation such as psychological stress, disease, accelerated growth, nutrient shortage and work load activate the HPA axis causing the release of CORT [23].

Thus, several hypothesized connections between CORT and telomere length exist. Firstly, CORT are an essential part of the vertebrate stress response, and their primary function is to mobilize energy [5]. Accordingly, the “metabolic telomere attrition hypothesis” proposes that during stress events that require an increased amount of energy and metabolic rates, telomeres are shortened as collateral [20]. As a result of the high energy expenditure, the energetically expensive maintenance of telomeres cannot take place as an emergency survival mechanism due to a shift in energy allocation [23]. In addition, CORT stimulate the generation of reactive oxygen species (ROS) and subsequent oxidative damage to telomeres, which are particularly susceptible to oxidation due to a high guanine content [11, 24–26]. Finally, cortisol reduces telomerase --the enzyme responsible for telomere maintenance-- activity in human T lymphocytes [27]. This reduction in telomerase activity can result in excessive telomere attrition [28]. Since wildlife face an array of stressors throughout their lifetime and these stressors can erode telomeres, CORT may play a mechanistic role in telomere loss [1].

External stressors cause pleiotropic effects that can potentially influence telomere dynamics, however the evidence for a causal relationship between CORT and telomere length is sparse. Thus, it is essential to build a quantitative understanding of the relationships between the HPA-mediated stress response and its downstream effects. In this study, we review the existing literature for empirical evidence of the relationship between CORT secretion and telomere length to better understand the underlying mechanism of telomere shortening as well as potential consequences of the neuroendocrine stress response in non-primate vertebrates. Using a meta-analytical framework, we tested whether CORT levels impact telomere length and if this relationship can differ among time frame, life history stage, or stressor type. We hypothesized that elevated CORT levels are linked to a decrease in telomere length.

## Methods

### Literature Search and Study Selection

We conducted a literature search for studies investigating the relationship between telomere length and CORT levels in non-human vertebrates using four search engines: Web of Science, Google Scholar, Pubmed and Scopus. Five subsets of the following keywords ‘reactive oxygen species,’ ‘antioxidant,’ ‘glucocorticoid,’ ‘cortisol,’ ‘corticosterone,’ ‘telomere length,’ ‘chronic stress,’ ‘oxidative stress,’ ‘acute stress,’ ‘chronic stress,’ ‘telomeres,’ and ‘HPA-axis,’ were conducted in each search engine. We did not specify a time frame in our literature search. Additional records were obtained from the reference section of studies included in the meta-analysis. Our study includes a qualitative synthesis of 31 full-text, peer-reviewed studies, and we report effect sizes for 15 of these studies.

Studies were excluded if (1) CORT was administered, but physiological measurements such as feather or plasma CORT levels were not taken. Such studies were excluded because it would not be possible to calculate the appropriate effect size (Fisher’s Z) for correlation data. For homogeneity in effect size calculation and statistical analysis, we did not include studies in which (2) CORT levels and telomere length measurements did not occur at the same or similar time points, (3) raw data was not accessible to use for the effect size calculation, or (4) telomere length measurements or CORT measurements were log transformed.

### Statistical Data Analyses

#### Meta Analysis

We conducted statistical analyses exclusively on studies with raw data available. When data was not publicly accessible, we contacted authors via email for consensual access. For each study, the correlation coefficient (R^2^) was calculated by fitting a linear mixed model using the “lme4” R package (version 3.6.1, R Development Core Team, Boston, MA). When possible, random effects such as multiple blood draws from a single individual were incorporated in the linear mixed model (LMER) to account for variability not captured by explanatory parameters. For studies where a random effect could not be determined, a linear model (LM) was fitted. From the LMs and LMERs, R^2^ values were obtained from the model and converted into Fisher’s Z, then adjusted for sample size and combined into a pooled effect size (Fisher’s Z; Z) using the R package “meta”. The random-effects model meta-analysis was implemented in our study as this model accounts for the assumption that studies come from different populations, rather than the same population. These pooled effect sizes were then visualized in a forest plot.

The “meta” package was also used to assess the statistical difference between observed and fixed effect model estimate of effect size (Cochrane’s Q) and the percent of variability in effect sizes that is not caused by sampling error (I^2^). After estimating heterogeneity, we identified potential outliers. Studies were classified as outliers if the study had an effect size with a confidence interval that did not intersect with the confidence interval of the pooled effect size.

Since some studies can have a larger influence on the pooled effect size than others due to its sample size or individual effect size, we conducted an influence analysis. The analysis was conducted by omitting each study one at a time and simulating the pooled effect size, with a confidence interval had the study not been included. This influence analysis was represented in a Baujat plot, which shows the contribution of each study to heterogeneity as Cochrane’s Q, and compares this to study’s influence on the pooled effect size.

#### Subgroup Analysis

Since experimental design can affect the outcome of a study, differences in effect size may be attributed to these variables. As such, further sources of between-study heterogeneity were investigated through subgroup analysis and meta-regression. In the subgroup analysis, studies were grouped based on different categorical experimental parameters. We completed five different subgroup analyses for the following parameters---duration of stressor, type of CORT assay, taxa, life history stage, and stressor type. For each subgroup analysis, a pooled effect size (Fisher’s Z) was calculated. We then compared pooled effect sizes and tested for between-study subgroup differences. The meta-regression was analogous to the subgroup analysis, except the parameter of investigation is continuous rather than categorical. We conducted one meta-regression for publication year and subsequently tested for between-study subgroup differences. For all analyses the significance threshold was set at p<0.05.

In the subgroup analysis, studies included in the meta-analysis were clustered based on categorical grouping and represented as a pooled effect size with a 95% confidence interval. The between study difference was indicated by Cochrane’s Q and the subsequent p-value for this statistical measure. The first subgroup analysis “stressor duration” organized studies based on the timeframe of the experiment--less than one week (n=1), one to two weeks (n=2), two to three weeks (n=7), three to four weeks (n=1), or longer than four weeks (n=3). The second subgroup analysis, “type of stress” compared anthropogenic (n=5), naturally occurring stress (n=6), or if stress was simulated by CORT administration (n=3). The subsequent subgroup analysis “life history stage, “differentiates studies based on pre-maturate study organisms (n=12), or post-maturate study organisms (n=2). Next, the subgroup “cortisol assay,” separates studies into those that quantified plasma cortisol (n=12) or non-plasma cortisol (n=2). Finally, the fifth subgroup analysis contrasts avian (n=12) and non-avian (n=2) studies.

#### Publication Bias

Published studies may not accurately represent the total studies investigating an area of research due to selective outcome reporting, missing studies and a higher likelihood of publication of studies reporting a significant (p<0.05) result. While proving selective outcome reporting and other forms of publication biases is challenging, missing studies can be visually represented using a funnel plot. Commonly, studies with small effect sizes and small sample studies are likely to be missing, which can be depicted with funnel asymmetry or holes in the funnel plot. We created a funnel plot by graphing effect size against study precision, defined as the standard error of the effect size to visualize potential publication bias. We also report Egger’s test, which is represented by the intercept, it’s confidence interval, and the associated p-value to determine if publication bias was statistically significant.

#### Risk of Bias in Included Studies

We assessed studies for missing outcome-level data, measurement of the outcomes, and outcome reporting in each included study. For the missing outcome-level data domain, we considered studies that could not report values for telomere length or CORT in less than 10% low risk. We designated studies that did not report these values for 10-50% of study organisms as moderate risk and studies that did not report values for over 50% of CORT or telomere length, as high risk. Secondly, we based risk of bias in the measurement of outcome on the type of CORT and telomere measurement. Low risk studies utilized plasma CORT or salivary CORT because these quantifications capture elevations related to a short-term stress event within minutes. Studies that measured CORT in fecal matter received a ranking of some concern because fecal CORT typically encapsulates cumulative stress over the day rather than CORT related to a particular environmental stressor. We considered studies that measured CORT in feathers high risk because feathers incorporate CORT in over one a month. Finally, for the risk of bias due to outcome reporting we denoted studies that based results off a subset of time points or measurements high risk. We denoted studies that report results based on all time points with low risk. We took these three domains into consideration when assessing overall risk of bias.

## Results

### Literature Search and Study Selection

We electronically screened 789 records for relevance from the following databases: Google Scholar (n=512), Web of Science (n=105), PubMed (n=72), and Scopus (n=100). 2113 additional records were hand screened from the reference section of the 31 studies used in qualitative analysis. Of the total 2902 records that were screened for relevance, 78 were removed as duplicates and 183 full-text articles were assessed for eligibility. Of the full-text articles, we removed 152 studies that did not fulfill our inclusion criteria. We statistically analyzed 15 of the remaining 31 studies, the ones that provided raw data for analysis [29–43]. The other 16 studies appeared to fit criteria but did not provide raw data for analysis [44–58]. The literature and study selection process is illustrated using a PRISMA diagram (Fig 1).

**Fig 1.**
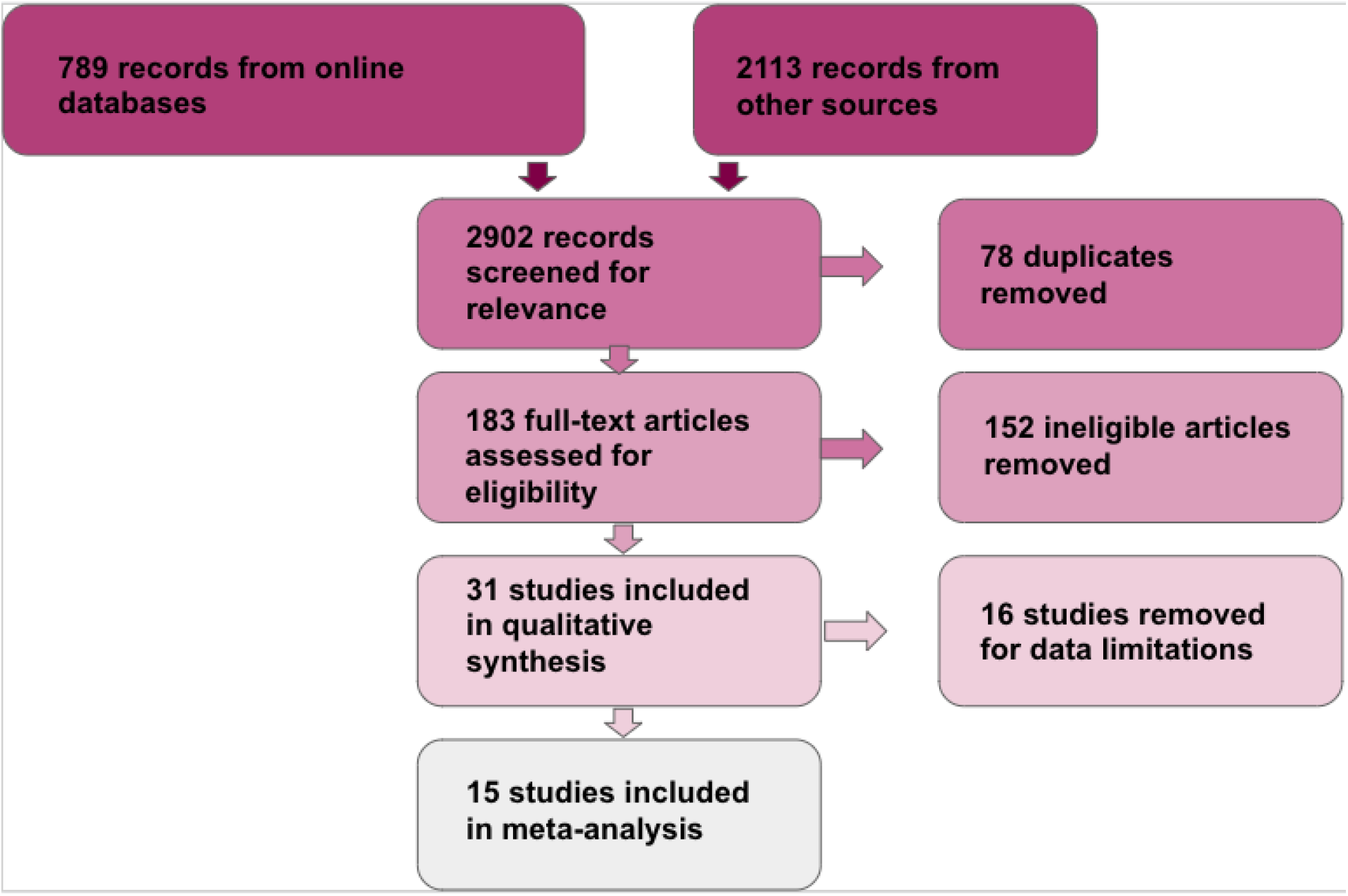
PRISMA diagram. PRISMA diagram showing the selection process for references included in the meta-analysis of the effects of CORT on telomere length.

### Meta-Analysis

The random-effects model meta-analysis is represented as a pooled effect size (Fisher’s Z) with 95% confidence intervals (Fig 2). No studies were removed as outliers. The model found no relationship between CORT levels and telomere length (Fisher’s Z= 0.1042, CI = 0.0235; 0.1836). Both heterogeneity measures, Cochrane’s Q (Q=11.31, p= 0.6615) and I^2^ with 95% confidence intervals (I^2^ = 0.0% ; CI = 0.0%; 42.6%) yielded similar results.

**Fig 2.**
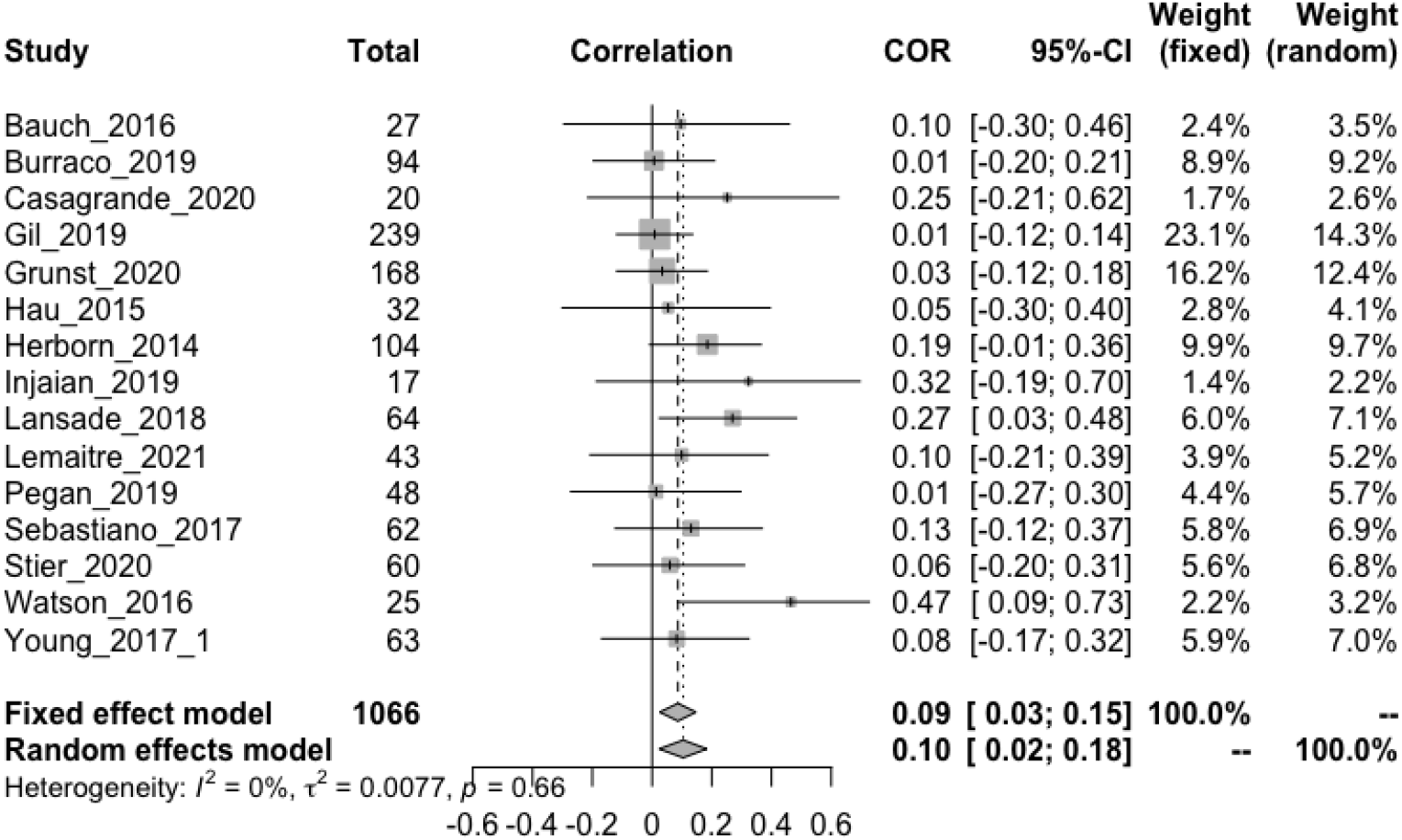
Forest plot. Distribution of effect sizes of CORT on telomere length and 95% CI of effect size. Dashed lines represent pooled effect sizes using a random and fixed effect model. Heterogeneity (I^2^), the percent of variability in effect sizes that is not caused by sampling error indicates very little variability in effect size. Weight indicates the influence the study has on the overall pooled effect.

The influence analysis indicated that theoretically removing one study at a time did not yield pooled effect sizes (Fisher’s Z=0.09-0.11) that differed greatly from the original pooled effect size (Fisher’s Z=0.11, Figure S3). Additionally, the influence analysis demonstrated that certain studies unevenly impacted the pooled effect size and/or overall heterogeneity (Figure S1), but no studies were removed as outliers.

### Subgroup Analysis

The subgroup analysis for “stressor duration” found no differences between any of the time frames (Table 1). The difference between-studies was not statistically significant (Q=1.86, p = 0.7594). Similarly, the subgroup analysis for the subgroup “stressor type,” did not reveal a difference between types of stressors (Table 1). The between study difference was not significantly different (Q=2.56, p=0.2783). Likewise, our subgroup “life history stage,” did not show differences between effect sizes for pre- and post-maturation organisms (Table 1), and did not indicate a difference between groups (Q = 0.06, p = 0.8119). The fourth subgroup analysis, “CORT assay” did not find a difference between plasma CORT and other CORT measurements, yielding a non-significant difference between studies (Q = 0.03, p = 0.8742) (Table 1). Lastly, our fifth subgroup analysis examined potential differences in effect size due to taxa, which could be divided into the binary categories avian and non-avian (Table 1). There was not a difference between-studies (Q = 0.03, p = 0.8666).

**Table 1.** Pooled effect sizes with 95% CI of experimental parameters investigated during the five subgroup analyses for stressor duration, stressor type, life history stage, cortisol assay and taxa group.

| Experimental Parameter | Number of Studies | Effect Size (Fisher's Z) | 95% CI |
| --- | --- | --- | --- |
| *Stressor Duration* |  |  |  |
| < 1 week | n=1 | 0.0135 | -0.2717; 0.2965 |
| 1-2 weeks | n=2 | 0.0959 | -0.1663; 0.3454 |
| 2-3 weeks | n=7 | 0.0741 | -0.0157; 0.1628 |
| 3-4 weeks | n=1 | 0.0957 | -0.2950; 0.4591 |
| > 4 weeks | n=3 | 0.2606 | 0.0075; 0.4823 |
| *Type of Stress* |  |  |  |
| Anthropogenic | n=5 | 0.1902 | 0.0059; 0.3621 |
| Naturally occurring | n=6 | 0.0410 | -0.0464; 0.1277 |
| CORT administration | n=3 | 0.1425 | -0.0320; 0.3085 |
| *Life History Stage* |  |  |  |
| Post-maturation | n=12 | 0.0729 | -0.1938; 0.3295 |
| Pre-maturation | n=2 | 0.1111 | 0.0185; 0.2019 |
| *Cortisol Assay* |  |  |  |
| Plasma Cortisol | n=12 | 0.1319 | -0.0941; 0.3449 |
| Non-Plasma Cortisol | n=2 | 0.1012 | 0.0061; 0.1945 |
| *Taxa Group*<br>Avian | n=12 | 0.1012 | 0.0088; 0.1919 |
| Non-Avian | n=2 | 0.1301 | -0.1248; 0.3688 |

The meta-regression was performed for the continuous variable publication year and represented as Cochrane’s Q and the associated p=value. Publication dates ranged from 2014-2021. Publication date was not a significant predictor of effect size (Q=1.252, p= 0.2632).

### Publication Bias

We found publication bias against studies with small sample size and small effect size (Figure S2; Egger’s test for small sample bias: intercept = 1.420616, CI= 0.3753223; 2.465909, p = 0.02064949).

### Risk of Bias in Included Studies

We represent the results of the risk of bias analysis in Table 2. Four of fifteen studies received a risk of bias ranking of moderate concern. These studies had some missing values for CORT or telomere length or selectively reported one time point in the results. The other eleven studies received a ranking of low risk and accordingly reported nearly all values for physiological parameters, measured CORT in plasma or saliva, and did not selectively report results.

**Table 2.** Overall risk of bias assessed based on missing outcome data, measure of outcome and in the selection of reported results.

| Author | Year | Bias due to missing outcome data | Bias in measure of outcome | Bias in the selection of reported result | Overall Risk of Bias |
| --- | --- | --- | --- | --- | --- |
| *Bauch et. al* | 2016 | high | low | high | *some concern* |
| *Burraco et. al* | 2019 | low | low | low | *low risk* |
| *Casagrande et. al* | 2020 | high | low | low | *some concern* |
| *Gil et. al* | 2019 | low | low | low | *low risk* |
| *Grunst et. al* | 2020 | low | high | low | *low risk* |
| *Hau et. al* | 2015 | low | low | low | *low risk* |
| *Herborn et. al* | 2014 | low | low | low | *low risk* |
| *Injaian et. al* | 2019 | high | low | high | *high risk* |
| *Lansade et. al* | 2018 | low | low | low | *low risk* |
| *Lemaitre et. al* | 2021 | low | moderate | low | *low risk* |
| *Pegan et. al* | 2019 | high | low | low | *some concern* |
| *Sebastiano et. al* | 2017 | low | low | low | *low risk* |
| *Stier et. al* | 2020 | low | low | low | *low risk* |
| *Watson et. al* | 2016 | low | low | low | *low risk* |
| *Young et. al* | 2017 | moderate | low | high | *some concern* |

## Discussion

External and internal stimuli can activate the neuroendocrine stress response in vertebrates, resulting in the secretion of CORT, which induces multiple downstream physiological and behavioral effects [8–9]. CORT might directly or indirectly cause telomere erosion [1, 11, 59]. Therefore, our goal was to investigate the relationship between CORT and telomere length *via* meta analysis using data from empirical studies. Though our sample size was limited (n=15), our data do not support the hypothesis that elevated CORT levels result in telomere shortening.

The empirical evidence for a relationship between CORT and telomere length is mixed, with some studies showing that telomere shortening is directly related to CORT levels, and other studies finding no relationship. For example, CORT influences telomere dynamics in wild roe deer and great tits [59–60], but not in red squirrels or magellanic penguins [61–62]. These results suggest that the relationship between CORT and telomere length is species-specific. Alternatively, a potential relationship may be obscured by the methods used to measure CORT and telomere length or by differences in experimental design including time frame. A differential sensitivity of the HPA-axis can also obscure conclusions made from CORT measurements especially in free-ranging vertebrates that can potentially encounter a variety of external stimuli [1]. For example, since CORT levels remain elevated for several minutes after a stressor subsides, it can be challenging to assess whether a measured cortisol increase results from the stressor in question, the stress involved in obtaining a sample from the experimental subject, or an unrelated event triggering HPA-axis activation [6, 63]. As baseline CORT samples must be taken quickly in many species, and in some cases less than three minutes, it can logistically be difficult to attain a true baseline CORT value in the field [64–67]. CORT can also be incorporated into other matrixes such as saliva, plasma, feathers, and hair [4, 65]. The multitude of non-invasive CORT sampling sources is advantageous to conservation physiology as their quantification does not require capture [6]. However, across tissues and fluids, the time required for CORT incorporation varies. For example, elevations in plasma cortisol can be detected within minutes of stressor exposure, whereas cortisol integrates into hair a week or more after an animal encounters a stressor [4]. Hence, there are caveats in the interpretation of each measurement such as incongruencies between plasma CORT and other tissues, hair and saliva [67]. Furthermore, CORT measurements in feces may be more representative of accumulated stress, rather than the event in question [6].

Similar considerations must be taken into account when assessing telomere length. Since telomere length can be influenced by environmental, maternal, and epigenetic effects, there is a large inter-individual variability in telomere dynamics [11, 68]. Several factors may contribute to this variability including discrepancies between the repeatability of different telomere measurement assays. Seven studies included in our meta-analysis utilized the telomere restriction fragment (TRF) assay, which depends on the distribution of the terminal restriction fragments to average the length of telomeres in a given cell population [69]. The other seven studies used the quantitative polymerase chain reaction (qPCR), which relies on the quantification of the highly conserved (TTAGGG)_n_ sequence for telomere quantification [70]. TRF-based studies are highly repeatable within individuals, whereas qPCR based studies are less repeatable and more variable than TRF because they are more prone to measurement errors [71]. qPCR can also bias measurements of telomere length because some species that exhibit interstitial telomeric repeats will artificially enlarge telomere length [72–73]. In addition to methodological differences, there is large individual variability in telomere length based on tissue type [74]. In adult zebra finches, telomere length in red blood cells is correlated with telomere length in the spleen, liver and brain, but not muscle or heart [74]. While avian studies in our meta-analysis used red blood cells for telomere measurement, telomere length was measured in tail muscle and liver in mammals and amphibians, which could lead to discrepancies when comparing among studies [61, 75–76].

A variety of biological factors also contribute to the diversity of telomere dynamics observed within a study and the large amount of observed inter-individual variability. The rate of telomere shortening can be influenced by the life histories and environmental conditions [22]. In accordance with the metabolic telomere attrition hypothesis, shortening is exacerbated by life history stages requiring more energy, such as reproduction [59]. Within an energy intensive process like reproduction, there can be a large inter-individual variability related to reproductive effort, which can be attributed to brood size and food availability [77]. Differences in reproductive roles during the breeding season account for sex-specific telomere dynamics which can contribute to differences in the variability of telomere dynamics within a study [78]. Finally, individuals respond differently to environmental challenges which can act synergistically with rapid growth or energy intensive life stages to magnify the rate of telomere shortening in non-model vertebrates [72].

Additionally, telomere dynamics can be complicated by the presence of telomerase which in some cases can elongate telomeres [79–80]. Typically, telomerase exhibits higher activity in developing organisms as compared to adults [81]. Ectotherms such as amphibians and reptiles have telomerase that is active throughout adulthood while endotherms reduce telomerase expression almost to non-detectable levels as they reach maturity [11, 71]. However, there is conflicting evidence on these observations, as telomerase activity has been detected in adult common terns and European Storm Petrels among other species [82–83]. Nonetheless, adult telomere shortening is observed in chickens, which have active telomerase in the adult life stage [49]. While there is an absence of empirical evidence on the long-term activity of telomerase in many avian species, even adults exhibit general shortening trends [80].

Many factors can potentially influence CORT and telomere measurements. During the subgroup analysis, we attempted to disentangle the underlying causes of the variation in effect size. Ultimately, we found no impact of stressor, taxa, type of CORT assay, or life history stage on the heterogeneity of the effect size. While no subgroup was identified as a predictor of heterogeneity in effect size, pooled effect sizes in certain categories with the subgroup indicate a higher pooled effect size than the overall pooled effect size. The small sample size for some parameters preclude further statistical analysis, however, we found variables of interest that may play a larger role into the relationship between CORT and telomere length. For example, within the subgroup “experimental timeframe,” (n=4) the group of studies with a timeframe above four weeks had a pooled effect size of 0.2181, while all other groups’ pooled effect size was less than that of the overall pooled effect size. Since most studies took place in less than four weeks, this suggests that while almost immediate changes in CORT can be observed, the impact of CORT on telomere length cannot be seen on short time scales. This idea is consistent with typical responses of telomere shortening observed in studies that take place for more than a year [83–87]. More work is needed to explore if long-term rather than short-term studies can be used to tease apart parameters that underlie the connection between CORT and telomere length such as stressor type or duration.

While CORT secretion is often viewed as the endpoint of HPA-axis activation in response to external stimuli, CORT manipulation is an oversimplification of the stress response that involves a multitude of physiological mechanisms that can each impact energy allocation and promote telomere erosion [8]. This highlights the problematic nature of the category “CORT stress” which was investigated as a category during the subgroup analysis, in which studies subjected organisms to CORT manipulation *via* an implant or oral administration. Since previous research found that organismal stress can result in adverse physiological responses without the involvement of the HPA-axis, these results highlight the issue of using CORT as a proxy for stress [88–89].

Overall, we found no relationship between CORT and telomere length across studies. Currently, the existing literature shows both a direct relationship and a lack of a relationship between CORT and telomere dynamics, suggesting that the underlying mechanisms driving this relationship are species-specific or altered by differences in experimental design. However, due to limited sample size, we are unable to investigate the underlying variables that play a role in this relationship. Here, we highlight the need for more studies with targeted experimental parameters to understand how conditions, such as experimental timeframes, stressor(s), and stressor magnitudes can drive a potential relationship between the neuroendocrine stress response and cellular aging. Thus, we recommend the following research priorities to groups studying similar questions.

1) Experimental timeframes and stressor magnitudes should be long enough to observe telomere erosion in relation to stressors when studying CORT.
2) When possible, studies should use a repeated measures design to measure cortisol levels and telomere lengths before and after stress exposure to account for individual variation.
3) While the avian taxa are well represented in this research topic, there is a dearth of information on other taxa. It will be important to investigate the neuroendocrine stress response of other vertebrates including mammals and reptiles to understand if similar principles hold true in these taxa or if telomere dynamics differ across taxa.

## Certainty of Evidence

We utilized the applicable Cochrane/GRADE categories “risk of bias,” “inconsistency,” and “publication bias,” for the determination of the certainty of evidence. Overall, we have a moderate confidence in the certainty of evidence. While most studies received a low risk of bias assessment, and had low heterogeneity, we report a considerable amount of publication bias as evidenced by Egger’s test and an asymmetrical funnel plot.

## Data Availability

The authors declare no competing interests. This study was registered in the Open Science Framework Registry (https://osf.io/rqve6). The review protocol can be accessed at https://bookdown.org/MathiasHarrer/Doing_Meta_Analysis_in_R/. Data are available from the Dryad Data Repository (https://datadryad.org/stash/dataset/doi:10.6078/D10T42) and will be made freely available upon acceptance.

## Acknowledgements

We would like to thank all the researchers who made their data freely available for this study. In particular, we thank Christine Bauch (University of Groningen), Stefania Casagrande (Max Planck Institute for Ornithology), Britt Heidinger (North Dakota State University), Marie-Pierre Moisan (French National Institute for Agriculture, Food, and Environment), Teresa Pegan (University of Michigan), Manrico Sebastiano (French National Centre for Scientific Research), Mathilde Tissier (Bishop’s University), and Hannah Watson (Lund University).

## Supporting information

**Figure S1.**
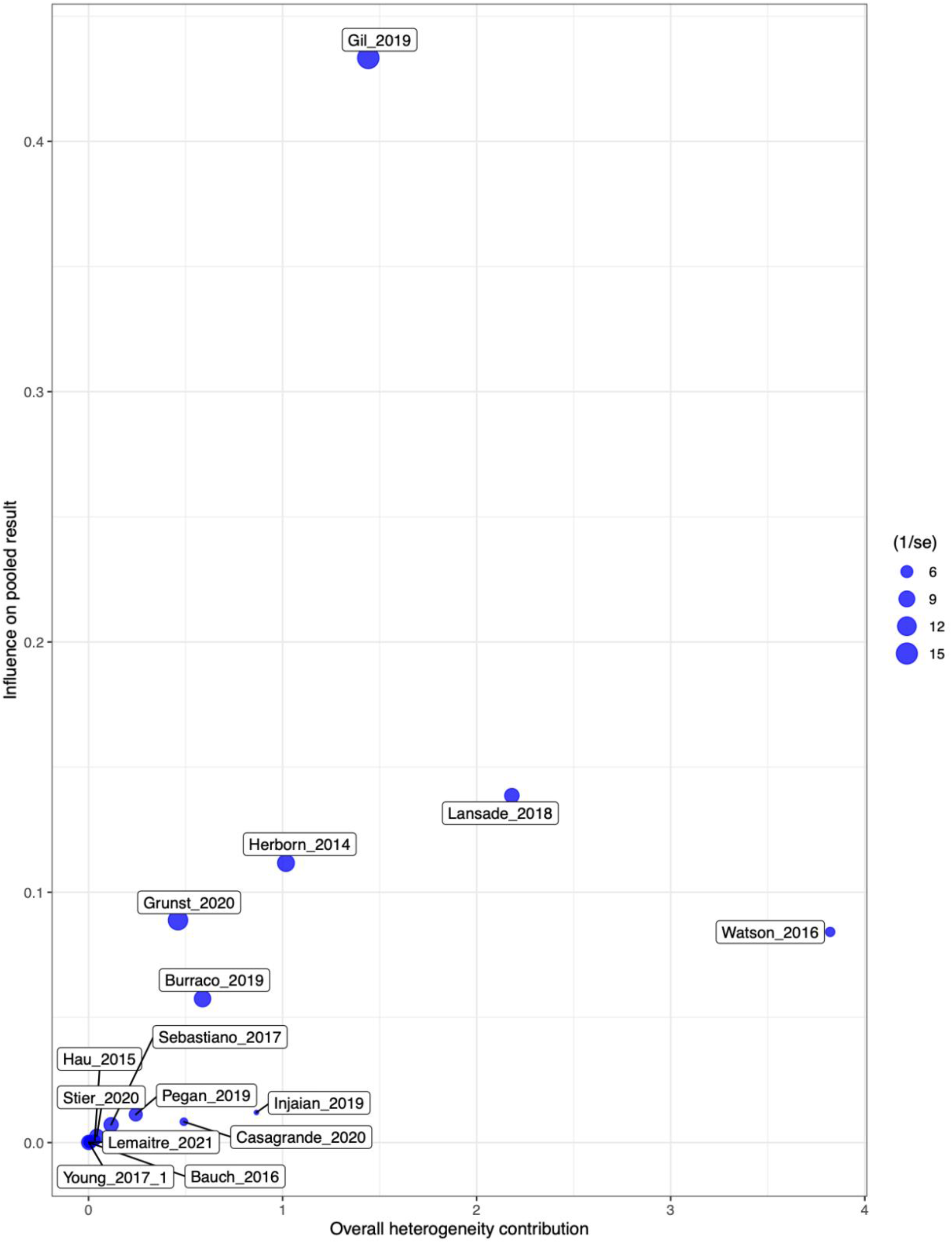
Baujat Plot. Studies can have an unequal influence on the pooled effect size and contribute to the heterogeneity of effect sizes. The horizontal axis represents Cochrane’s Q and influence on the pooled effect size on the vertical axis.

**Figure S2.**
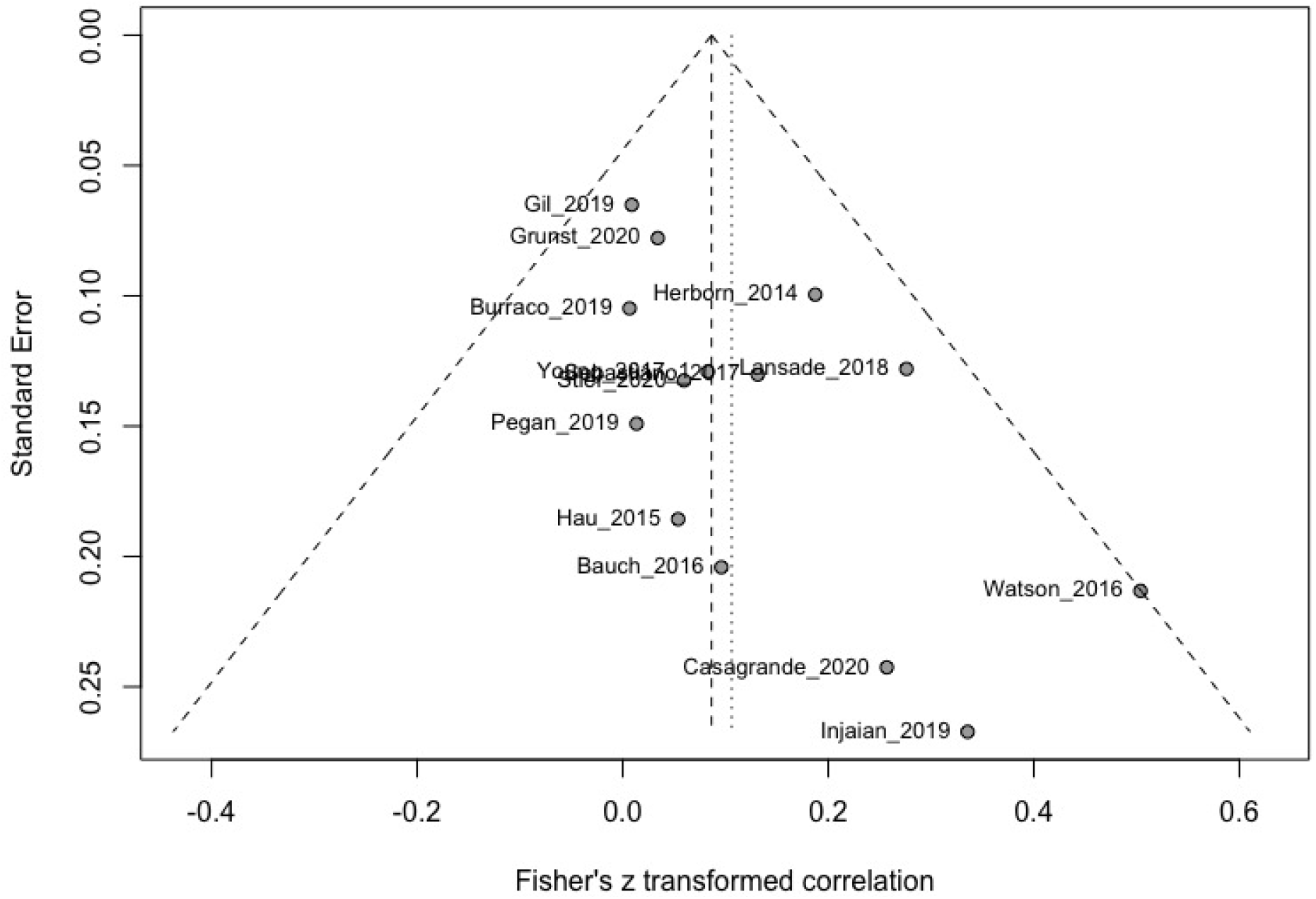
Funnel Plot. The lack of studies in the bottom left of the “funnel” demonstrates publication bias against studies with small sample sizes and small effect sizes.

**Figure S3.**
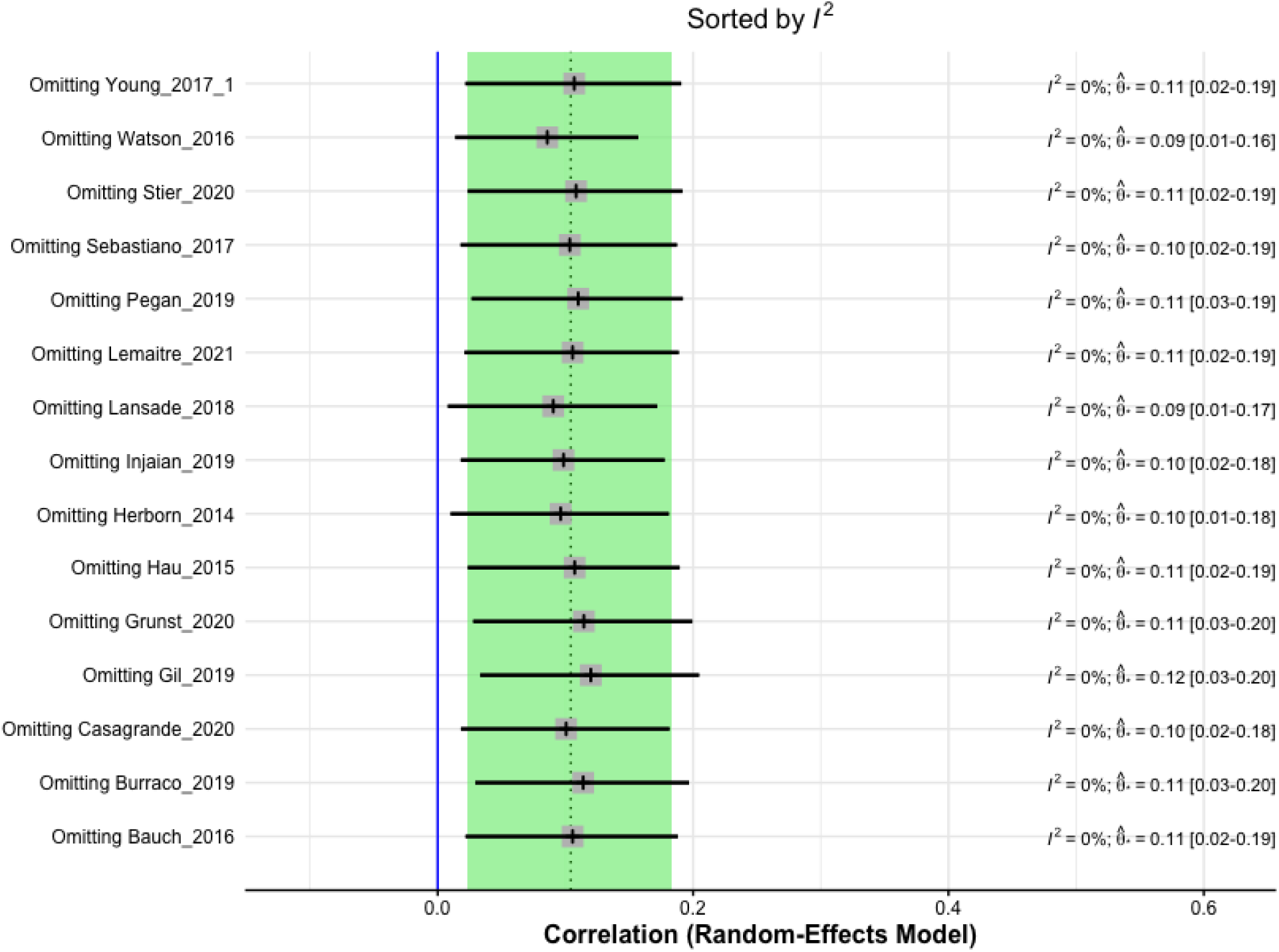
Influence Analysis Plot. The leave one out recalculation reveals a similar effect size across studies and indicates that studies evenly contribute to the pooled effect size.

**Table S1.** Search Strategy Table. Details search term combinations used to search online databases and websites.

